# Affinity tag free purification of SARS-Cov-2 N protein and its crystal structure in complex with ssDNA

**DOI:** 10.1101/2024.09.16.613250

**Authors:** Atanu Maiti, Hiroshi Matsuo

**Author notes:** To whom correspondence should be addressed. Atanu Maiti, Ph.D., Hiroshi Matsuo, Ph.D.

## Abstract

The nucleocapsid (N) protein is one of the four structural proteins in SARS-CoV-2, playing key roles in viral assembly, immune evasion, and stability. One of its primary functions is to protect viral RNA by forming the nucleocapsid. However, the precise mechanisms of how the N protein interacts with viral RNA and assembles into a nucleocapsid remain unclear. Compared to other SARS-CoV-2 components, the N protein has several advantages: higher sequence conservation, lower mutation rates, and stronger immunogenicity, making it an attractive target for antiviral drug development and diagnostics. Therefore, a detailed understanding of the N protein’s structure is essential for deciphering its role in viral assembly and for developing effective therapeutics. In this study, we report the expression and purification of a soluble recombinant N protein, along with a 1.55Å resolution crystal structure of its nucleic acid-binding domain (N-NTD) in complex with ssDNA. Our structure reveals new insights into the conformation and interaction of the flexible N-arm, which could aid in understanding nucleocapsid assembly. Additionally, we identify residues that are critical for ssDNA interaction.

## 1. Introduction

The Covid-19 pandemic, caused by Severe Acute Respiratory Syndrome Coronavirus – 2 (SARS-Cov-2) is one of the most devastating health and socio-economic crisis in modern history [1–3]. According to World Health Organization (WHO) about 7.1 million people have died and 776 million people have infected by SARS-COV-2 infection till date (https://data.who.int/dashboards/covid19/cases). Although, timely development and world wise implementation of vaccines have paused the pandemic, the newly emerging variants still poses threat to human health and for future pandemic [4,5]. Therefore, effort to better understand SARS-CoV-2 and development of antiviral agents are ongoing since after the pandemic started [6–8].

SARS COV-2 is a positive-sense single-stranded RNA (ssRNA) virus, belongs to beta-coronavirus family and share genetic and structural homology with SARS-CoV (caused 2002 SARS epidemic) and MERS-CoV (emerged during 2012) [9,10]. The large RNA genome (about 30kb) of SARS COV-2 encodes 16 non-structural proteins (NSPS), at least 6 accessory proteins and 4 structural proteins, namely Spike (S), Nucleocapsid (N), Membrane (M) and Envelope (E) [11] (Fig 1a). These proteins play important role in viral life cycle, host infection and pathogenicity [12,13]. While, S, M and E proteins mainly engage in the outer shell of the virus particle, N protein comprises viral capsid by packaging the viral RNA genome into helical ribonucleoprotein (RNP) complex [13,14]. Thus, N protein plays a pivotal role in protecting viral genome forming nucleocapsid inside viral particle and less prone to mutation [15,16]. N protein also participates in other processes like viral genome transcription, replication and immune regulation by interacting with viral and host proteins. N protein is highly immunogenic, most abundant viral protein during infection and highly conserved among other coronaviruses. These advantages make N protein a potential target for antiviral drug development and diagnosis and hence an attractive subject for structural studies [12,17–23].

**Figure 1:**
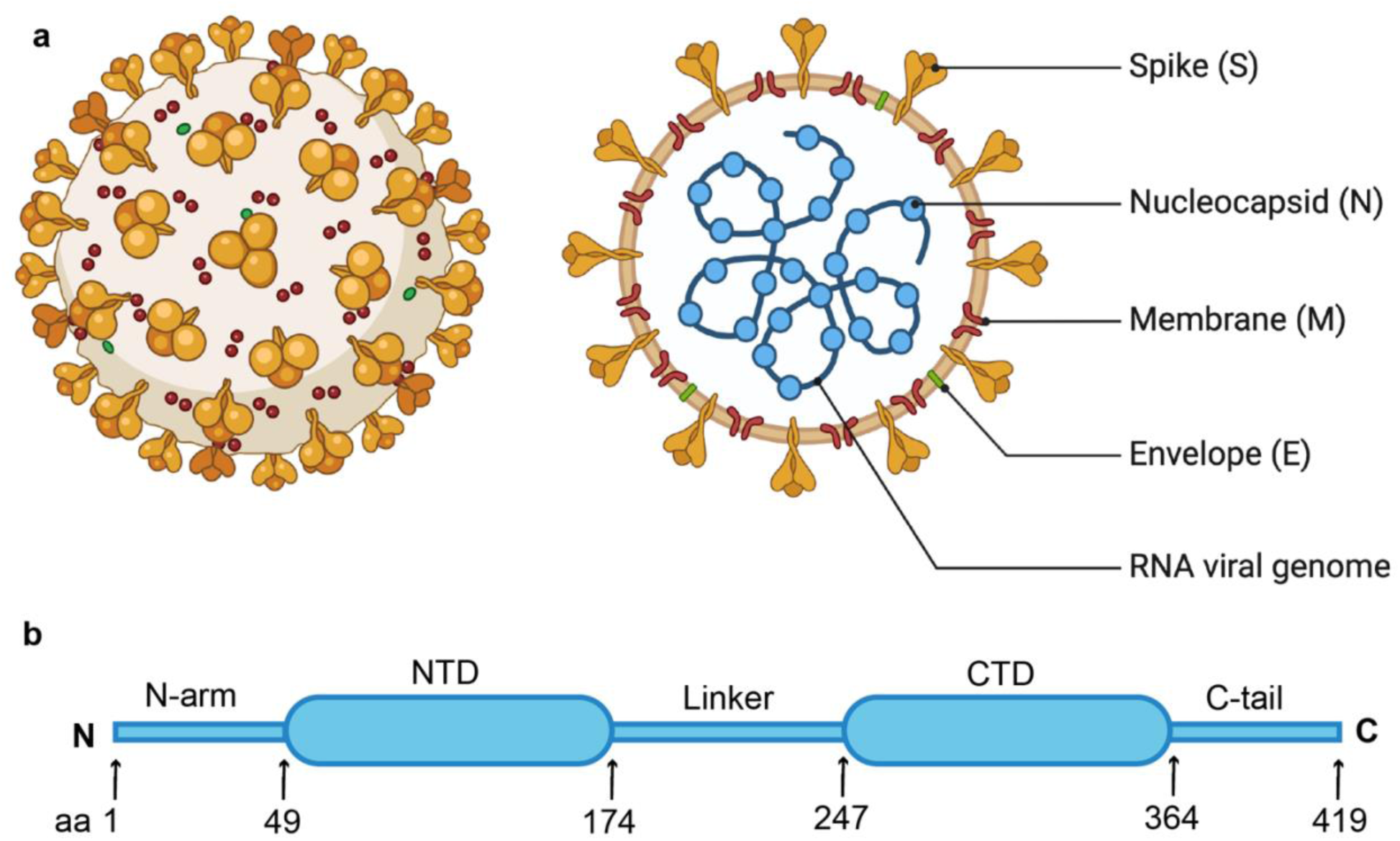
a) Schematic representation of SARS-CoV-2 virus and its structural proteins (S, N, M and E) with genomic RNA. b) The N protein is composed of several distinct regions; the N-terminal domain (NTD) and the C-terminal domain (CTD) connected by a flexible linker, and intrinsically disordered regions like the N-arm and the C-tail.

Numerous efforts have been given to understand the structure and function of coronaviruses N proteins. Structures of SARS-Cov-2 N protein consists of 419 amino acids comprising two structural domains, namely N-terminal domain (N-NTD; residues 48-174) and C-terminal domain (N-CTD; residues 247-364). These two domains are connected by a highly flexible linker region (residues 175-246) and other ends of these domains holds intrinsically disordered regions, namely N-arm (at N-terminal; residues 1–47) and C-tail (at C-terminal; residues 365–419) [24] (Fig1b). While N protein interacts with RNA through N-NTD, N-CTD is involved in self-dimerization and RNA binding [25–32]. The flexible linker region (residues 175-246) regulates the interaction of the N protein with RNA and other viral and host proteins [33,34]. Though it is believed that the disordered C-tail plays role in interaction with viral M protein and packaging signal [22,35], the role of N-arm is still elusive. Many structures of SARS-Cov-2 N protein N-NTD and N-CTD have been solved separately [23,25,28–32,36–44]. Recently, structures of SARS-Cov-2 full-length N protein have been solved by Cryo-EM at 6.00 Å and 4.57 Å resolution as dimer and monomer respectively [45]. However, high resolution structure of SARS-Cov-2 full-length N protein still in demand.

Escalating research on N protein need isolation and purification of high-quality N protein at large extent, which generally achieve by over expressing recombinant protein in bacterial cell. Several groups have been reported the purification of recombinant N protein from E. Coli using affinity tag [31,46–49]. Although affinity tag eases the protein purification procedure, retention or incomplete removal of affinity tag are not always ideal for downstream studies with the protein [50–55]. Moreover, the method used to purify can also impact the quality and molecular properties of N protein [56]. Therefore, developing a procedure to purify high quality SARS-Cov-2 full-length N protein without any affinity tag is worthy.

Here we report an affinity tag free purification method for recombinant full-length N protein of SARS-Cov-2 at large quantity and high purity. Our purified N protein is highly soluble, homogeneous, nucleic acids free and suitable for biochemical, biophysical and structural studies. To advance our understanding on the nucleocapsid assembly formation and structure, we aimed to crystalize full-length N protein in complex with nucleic acids (ssDNA). Our crystallization effort, resulted a high resolution (1.55 Å) structure of truncated full-length N protein, showing N-terminal domain (N-NTD) with part of flexible N-arm in complex with ssDNA. Crystal packing of our structure indicate a possible role of flexible N-arm in RNP assembly formation. Our structure shows ssDNA binds to N-NTD at the same RNA binding pocket, but the mode of interaction is different from RNA interaction. Our structure also reveals the amino acids residues important for ssDNA binding. As inhibition of nucleocapsid assembly is a major path for antiviral development, our structure could be beneficial to structure guided design of inhibitors.

## 2. Results and Discussion

### 2.1. N Protein production

Our goal was to purify SARS-CoV-2 N protein (full-length) to study the condition of its assembly formation and its structure in complex with nucleic acids. We developed expression vector for SARS-Cov-2 N gene (Gene ID: 43740575, codon optimized for E. Coli) without any affinity tag cloned in pET-11a vector (GenScript) and transformed the expression plasmid in E. Coli BL21 (DE3) cells to overexpress the protein. Cells were grown at 37°C till optical density 0.5-0.6 at 600 nm and were induced with IPTG at 17°C and continued to grow for overnight to express soluble protein at high concentration. Since N protein has high affinity with nucleic acids, we used polyethyleneimine precipitation method to separate nucleic acids and proteins. At final polyethyleneimine concentration of 0.5%, N protein remained in solution and was separated by centrifugation. The treatment of supernatant with SP Sepharose and elution with high salt (0.4-0.5M NaCl) allowed us to purify N protein to significant purity. Finally, eluted N protein was further purified by running through the Superdex-75 size exclusion column. The protein eluted was a single major species with high purity as appeared by size exclusion chromatograph and SDS-PAGE analysis (Fig 2a). Analysis of UV absorption spectra at 280 nm and 260 nm showed the absence of nucleic acids contamination in purified N protein. As reported by other groups [16,22] our mass photometry (MP) analysis showed that N protein remained as a homogeneous dimer at 50 to 100 nM protein concentration (Fig 2b). Overall, our purified N protein was highly soluble, homogeneous, nucleic acids free and suitable for biochemical, biophysical and structural studies.

**Figure 2:**
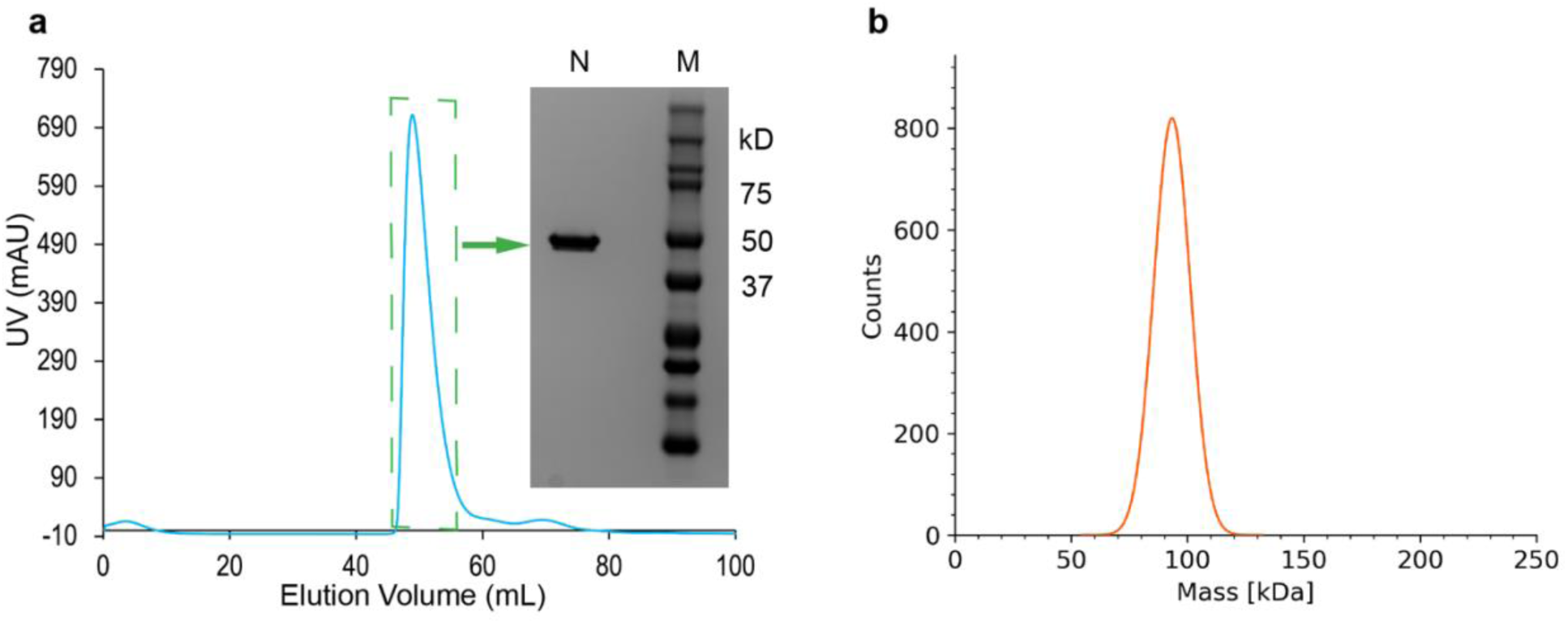
SARS-CoV-2 N protein was purified by Polyethyleneimine precipitation followed by SP Sepharose ion-exchange and finally size-exclusion chromatography. a) FPLC size-exclusion chromatograph and SDS-PAGE electrophoresis of purified N protein. N represents purified full-length N protein and M represents protein marker. b) Mass Photometry analysis revealed N protein appeared as a monodisperse dimer (∼93 kD) in solution.

### 2.2. Crystal formation and Structure of N-NTD with part of flexible N-arm

We intended to capture the oligomeric structure of full-length N protein in complex with nucleic acids. As reported by Zhao et al [57], trend of dimeric N protein’s higher order oligomer formation is dependent on nucleic acids length and showed strong binding affinity with 10nt poly-T (T-10) and 20nt poly-T (T-20) forming higher oligomer. Accordingly, we decided to crystalize N protein in complex with T-10 (10nt T) and T-20 (20nt T). Towards our aim, we set up different crystallization conditions for N protein in complex with T-10 (10nt T) and T-20 (20nt T) simultaneously at two different temperatures 4°C and 20°C using different commercially available crystallization screens. After an extended period, crystals appeared in a single condition (Hampton Research, Peg/pH, F4) at 4°C, in a well having N protein and T-20 (Supplementary Fig. S1a). Crystal cryoprotected with 20% glycerol, was diffracted at high resolution at 22-ID beam line, APS synchrotron facility (Supplementary Fig. S1b). After diffraction data processing and analysis, the crystal appeared in the I 41 space group with unit cell parameter (a=90.891, b=90.891, c=36.341, α=90, β=90, γ=90) (Table-1). The cell content analysis in CCP4 program suit suggested that the unit cell volume cannot accommodate full-length N protein however can accommodate half of the full-length N protein. Molecular replacement with available structure coordinate of N-NTD (PDB ID:7N0R) and N-CTD (PDB ID:6ZCO) together (For full-length protein) or with only N-CTD (for C-terminal domain) failed to give any solution. However, molecular replacement with only N-NTD (for N-terminal domain) successfully provided a solution, resulting a single protein molecule per asymmetric unit and final structure determination at 1.55 Å resolution. Our finding of crystallization of only N-terminal part of the full-length protein after an extended period, made us to think that the full-length protein might had digested in the crystallization condition and then only N-terminal part had crystallized in the given condition. To verify our thought, we collected the residual from the corresponding crystallization wells of both with T-10 and T-20 and ran in SDS-PAGE gel electrophoresis. The SDS-PAGE confirmed that the full-length protein was indeed digested in parts under the crystallization condition producing a significant part with approximate molecular weight of 20 KD (Supplementary Fig. S1d & S1e). Supporting previous reports, our findings indicate that crystallization of full-length N protein is challenging due to its intrinsic flexibility and protease sensitivity [16,23,58,59]. Although we were uncertain about the exact length of the crystallized digested part of the protein, we were able to build the structure model according to the electron density observed after phase determination using molecular replacement. As usual, strong electron density observed for N-NTD structured region (residues 49-173) along with a part of flexible N-arm (residues 39-48) allowed us to build the structure model confidently. We were unable to build the model for rest of the protein due to lack of electron density. Overall, we solved the crystal structure of N protein from residues 39 to 173, part of flexible N-arm with entire N-NTD (Supplementary Fig. S1c & S1f and Fig. 3a). We also observed strong electron density for a nucleotide with additional strong density for two adjoining phosphates and poor density for corresponding base. As our crystallization sample contained 20nt T, the electron density observed here could be a stable part of the poly-T interacting with N-NTD and rest of the poly-T is not visible as flexible part. So, we modeled the trinucleotide as TTT (Supplementary Fig. 1c and Fig. 3a).

**Table-1:**
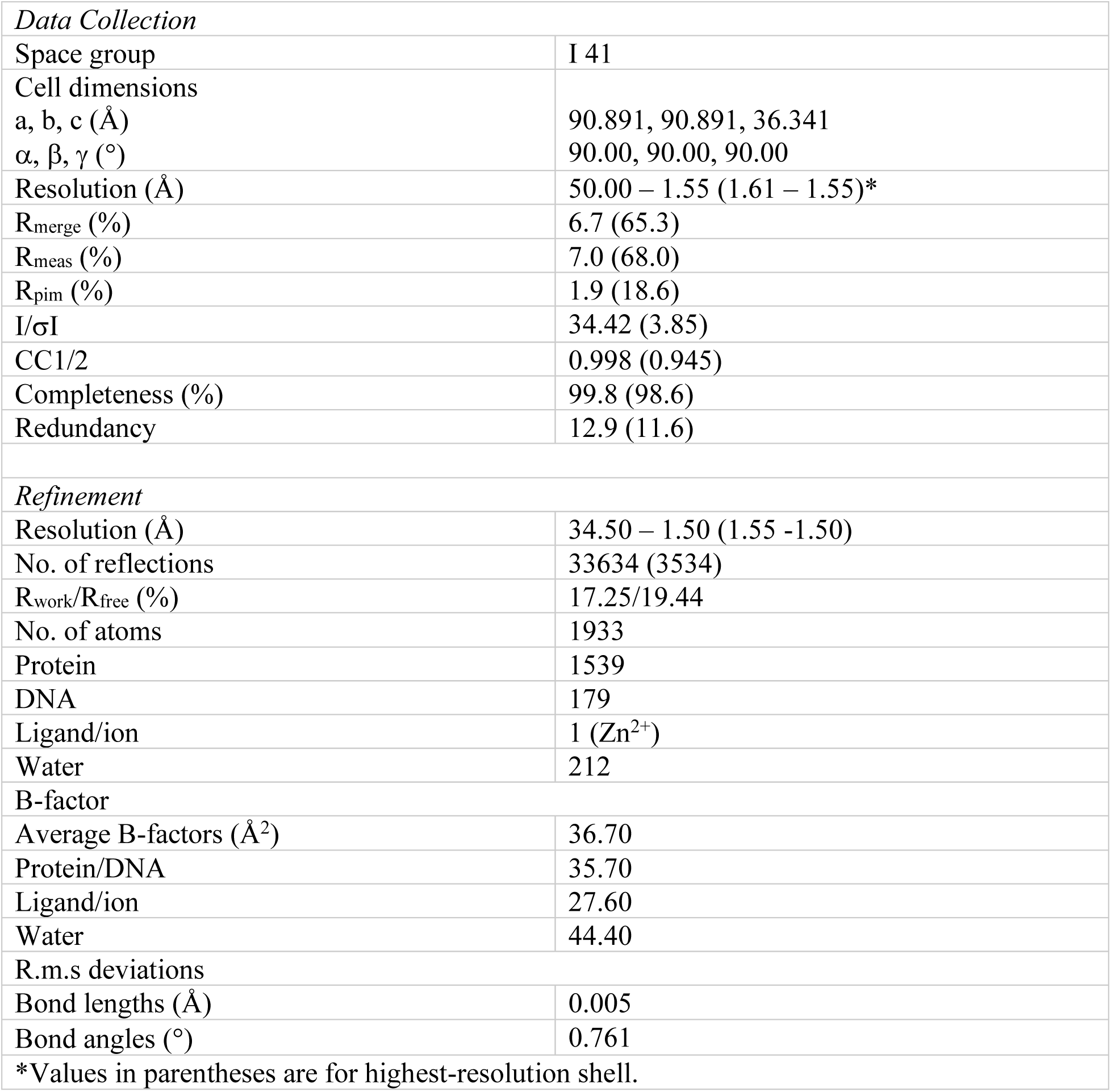
Data collection and refinement statistics.

|  |  |
| --- | --- |
| *Data Collection* |  |
| Space group | I 41 |
| Cell dimensions |  |
| a, b, c (Å) | 90.891, 90.891, 36.341 |
| $\alpha$, $\beta$, $\gamma$ (°) | 90.00, 90.00, 90.00 |
| Resolution (Å) | 50.00 – 1.55 (1.61 – 1.55)* |
| R<sub>merge</sub> (%) | 6.7 (65.3) |
| R<sub>meas</sub> (%) | 7.0 (68.0) |
| R<sub>pim</sub> (%) | 1.9 (18.6) |
| I/σI | 34.42 (3.85) |
| CC1/2 | 0.998 (0.945) |
| Completeness (%) | 99.8 (98.6) |
| Redundancy | 12.9 (11.6) |
| *Refinement* |  |
| Resolution (Å) | 34.50 – 1.50 (1.55 -1.50) |
| No. of reflections | 33634 (3534) |
| R<sub>work</sub>/R<sub>free</sub> (%) | 17.25/19.44 |
| No. of atoms | 1933 |
| Protein | 1539 |
| DNA | 179 |
| Ligand/ion | 1 (Zn<sup>2+</sup>) |
| Water | 212 |
| B-factor |  |
| Average B-factors (Å<sup>2</sup>) | 36.70 |
| Protein/DNA | 35.70 |
| Ligand/ion | 27.60 |
| Water | 44.40 |
| R.m.s deviations |  |
| Bond lengths (Å) | 0.005 |
| Bond angles (°) | 0.761 |
| *Values in parentheses are for highest-resolution shell. |  |

**Figure 3:**
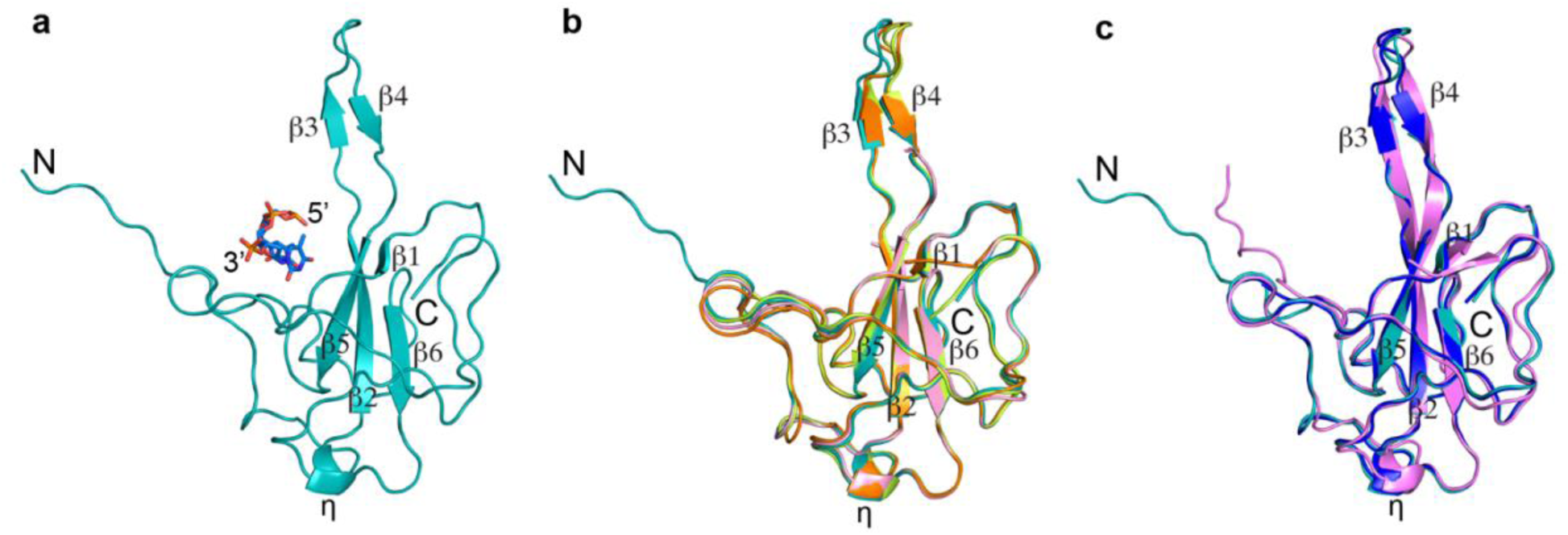
Comparisons of Coronavirus N-protein structures. a) Our N-protein structure (digested full-length N) showing N-NTD with part of flexible N-arm (teal color, cartoon) in complex with ssDNA (blue, stick). b) Superimposition of our structure with other published SARS-CoV-2 N-NTD structures (PDB ID:7XWZ; light pink color, PDB ID:7CDZ; orange color, PDB ID:7N0R; limon color). c) Superimposition of our structure with closely related SARS-CoV N-NTD (PDB ID: 2OFZ; navy blue color) and MERS-CoV-N-NTD (PDB ID: 4UD1; purple color) structures. N and C indicate the N- and C-terminal ends of the protein respectively. 5ʹ and 3ʹ represent 5ʹ-end and 3ʹ-end of ssDNA. ß and η represent ß-sheets and short 3_10_ helices respectively.

### 2.3. Comparison with published N-NTD structures

Since the covid-19 pandemic, several structures of SARS-CoV-2 N-NTD have been published: Some in complex with dsRNA or ssRNA [28,30], some with antibody/protein [37,38,42,44], some with small molecules and some as apo form [23,25,29,30,44]. Among all the crystal structures deposited in PDB, apo N-NTD crystallized as trimer (PDB ID: 6VYO, 6WKP, 7UW3, 6M3M, 7CDZ, 7XX1 and 7VBD); N-NTD in complex with dsRNA crystallized as a dimer (PDB ID: 7XWZ) and N-NTD in complex with antibodies or protein crystallize as monomer (PDB ID: 7STR,7STS,7SUE,7N3C,7N3D, 7CR5,7N0R,7R98 and 7VNU) in an asymmetric unit. Our N-NTD in complex with ssDNA crystalized as a monomer in an asymmetric unit (Fig. 3a) and maintains good topological agreement with other published crystal structure of SARS-CoV-2 N-NTD (Fig. 3b) as well as with closely related SARS-CoV (PDB ID: 2OFZ) and MERS-Cov N-NTD (PDB ID: 4UD1) structures (Fig. 3c). Likewise, N-NTD shaped like a right-handed fist, containing four-stranded antiparallel ß-sheets (ß1- ß5- ß2- ß6) core, sandwiched between two loops, a short 3_10_ helices (η) and a protruding ß-hairpin (ß3 and ß4) between ß2 and ß5 (Fig.3a-c). As previously reported, the protruding ß-hairpin (ß3 and ß4) region is flexible ranging a dynamic conformation in different structures. Previous N-NTD structures were mostly determined using the stable N-NTD construct residue 48 to174. However, in our structure (digested full-length N), in addition to the stable N-NTD, we determined structure of a part of flexible N-arm at high resolution. N-arm goes outward to the N-NTD core as observed in MERS-Cov N-NTD structure (Fig. 3c) [60].

### 2.4. Putative role of N-arm in Nucleocapsid assembly

In viral life cycle, the main role of N protein is to bind viral genomic RNA to form viral nucleoprotein complex (RNP) and assembling the RNP inside the virus particle. Previous studies pointed out that assembly of RNP complex involves multiple regions of N proteins that mediate protein-RNA and protein-protein interactions [17,22,61–63]. However, the mechanism of RNP assembly formation is still elusive. It is known that intrinsically disordered proteins (IDPs) or intrinsically disordered regions (IDRs) of a protein lack a defined structure, but play important roles in biological processes, particularly in macromolecular interactions [64–67]. Due to its intrinsic disorder nature, the structural information as well as biological role of N-arm of SARS-CoV-2 N protein is not clear yet. Our crystal structure shows, N-NTD with a part of flexible N-arm arrange in a specific pattern where N-arm extended to the neighboring molecules and anchored between two molecules through significant hydrogen bonding (Fig. 4). The mainchain amine of L45 form direct hydrogen bond with sidechain carbonyl of Q160. Mainchain amine and carbonyl of Q43 form direct hydrogen bond with sidechain carbonyl of N75 and mainchain amine of T76 respectively. Mainchain carbonyl of R41 form direct hydrogen bond with sidechain amine of N77. Sidechain N of R41 form a direct hydrogen bond with side chain NH_2_ of R93 of another neighboring molecule. Crystal packing of our structure suggest a possible role of flexible N-arm in RNP assembly formation. However, additional experiment will need to support this hypothesis.

**Figure 4:**
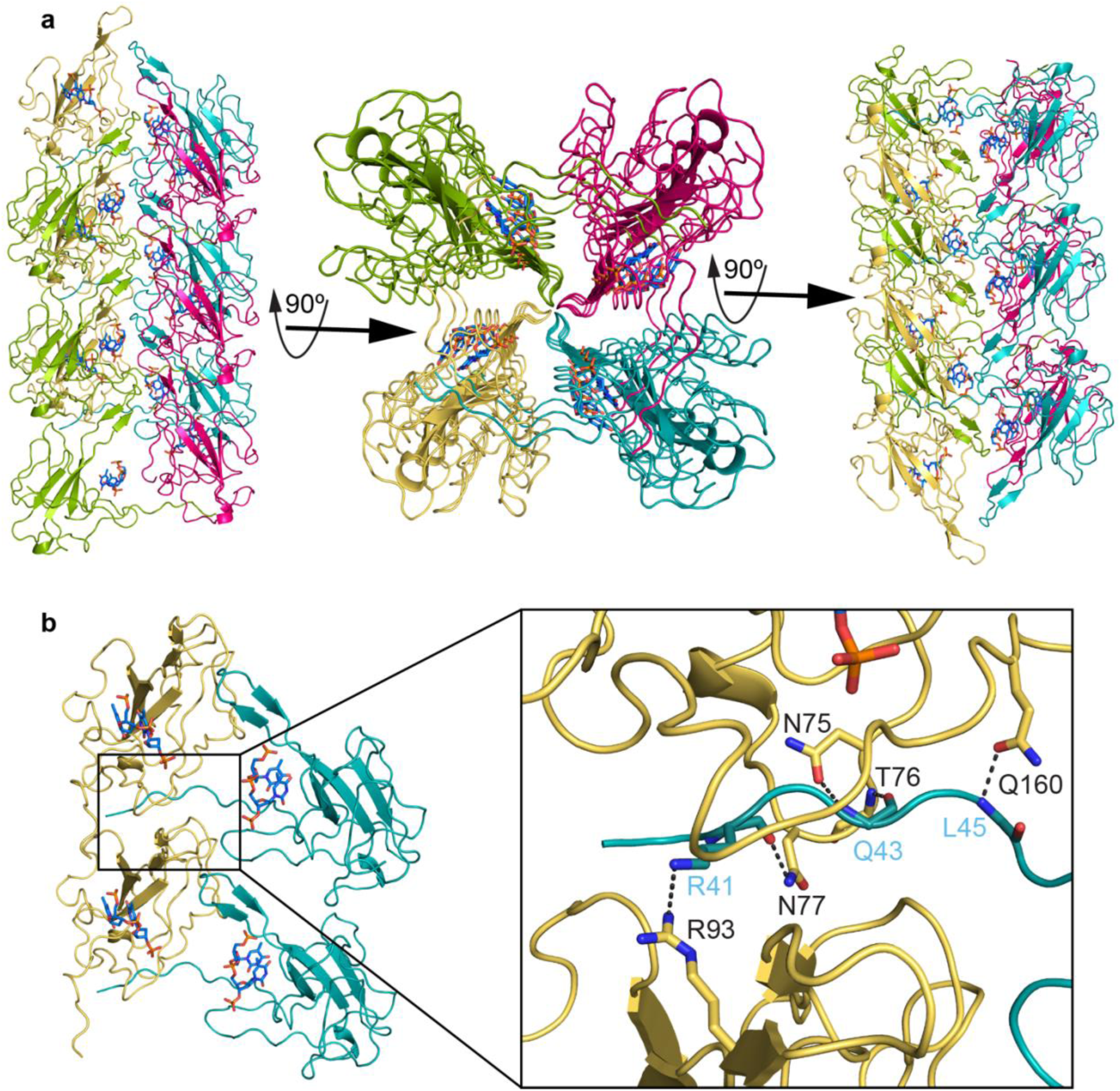
Crystal packaging of digested N protein. a) In crystal, four neighboring protein molecules are colored as teal, hotpink, splitpea and yelloworange showing crystal packing at different 90° angles. b) N-arm extends to the neighboring molecules and is anchored between two molecules through hydrogen bonding. Gray dashed lines represent the hydrogen bonds between amino acids.

### 2.5. ssDNA interacts at the RNA binding pocket of N-NTD with different interaction mode

Despite the main role of N protein as viral RNA genome packaging, new findings emphasize several biological processes which require interaction of N protein with ssDNA or dsDNA [68–70]. Previous computational and structural approaches provided significant information of how N protein interacts with dsRNA and ssRNA [28,30]. However, structural information of how N protein interacts with dsDNA or ssDNA are absent. To fill this gap, we aimed to crystalize poly - T (T-20) in complex with N protein. In our structure, we observed well define electron density for one T and additional strong density for two adjoining phosphates and partial density for corresponding base. The absence of electron density for other part of nucleotide could be due to the dynamic nature of ssDNA or ssDNA has been digested under crystallization condition [71–73]. As we were not sure about the corresponding sequential number of T in the structure, we numbered the T with well-defined electron density as 0 and -1 and +1 towards 5′ and 3′ respectively. Nevertheless, our structure provides some valuable information of N protein interaction with ssDNA. Superimposition of our structure with previous structures of N-NTD in complex with dsRNA and ssRNA (PDB ID: 7XWZ and 7ACT) shows that ssDNA binds to N-NTD at the same RNA binding pocket but in different mode (Fig. 5a & 5b). The Watson-Crick face of ssDNA interact with protein through direct and water mediated hydrogen bonds as well as π–π stacking. The Watson–Crick face of T_0_ has two direct and one water mediated interactions with the protein. The 2-carbonyl group forms a hydrogen bond with the mainchain amino proton of S51, N3 atom forms a hydrogen bond with the sidechain -OH group of Y111, and the 4-carbonyl group forms a water mediated hydrogen bond with the guanidino group of R88. T_0_ also stack with Y109 through π–π stacking interaction. Furthermore, the deoxy-ribose O3ʹ of T_0_ form direct hydrogen bond with the guanidino group of R149 (Fig. 5c). The Watson–Crick face of T_-1_ interacts with protein through one direct and one water mediated hydrogen bond. The 2-carbonyl group forms a hydrogen bond with the sidechain amino group of N47 and N3 atom forms a water mediated hydrogen bond with mainchain carbonyl group of T49. Additionally, the 5ʹ-phosphate group of T_+1_ is supported by one direct hydrogen bond with the guanidino group of R149 and one water-mediated hydrogen bonds with the mainchain carbonyl group of P151(Fig. 5c). On the other hand, RNA interacts with N protein through phosphate back bone (Fig. 5d) [28,30]. Residues R88, Y111, Y109 and R149 are involved in interaction with both DNA as well as RNA. Recent studies highlighted the use of oligonucleotide as potential inhibitors of protein function [74]. A designed ssDNA aptamer has shown promises as an antiviral therapy against Covid-19 which disrupt the liquid-liquid phase separation (LLPS) mediated by the N protein [70]. Our structure along with previous structures will aid to design better nucleic acids base therapeutics against SARS-Cov-2 virus.

**Figure 5:**
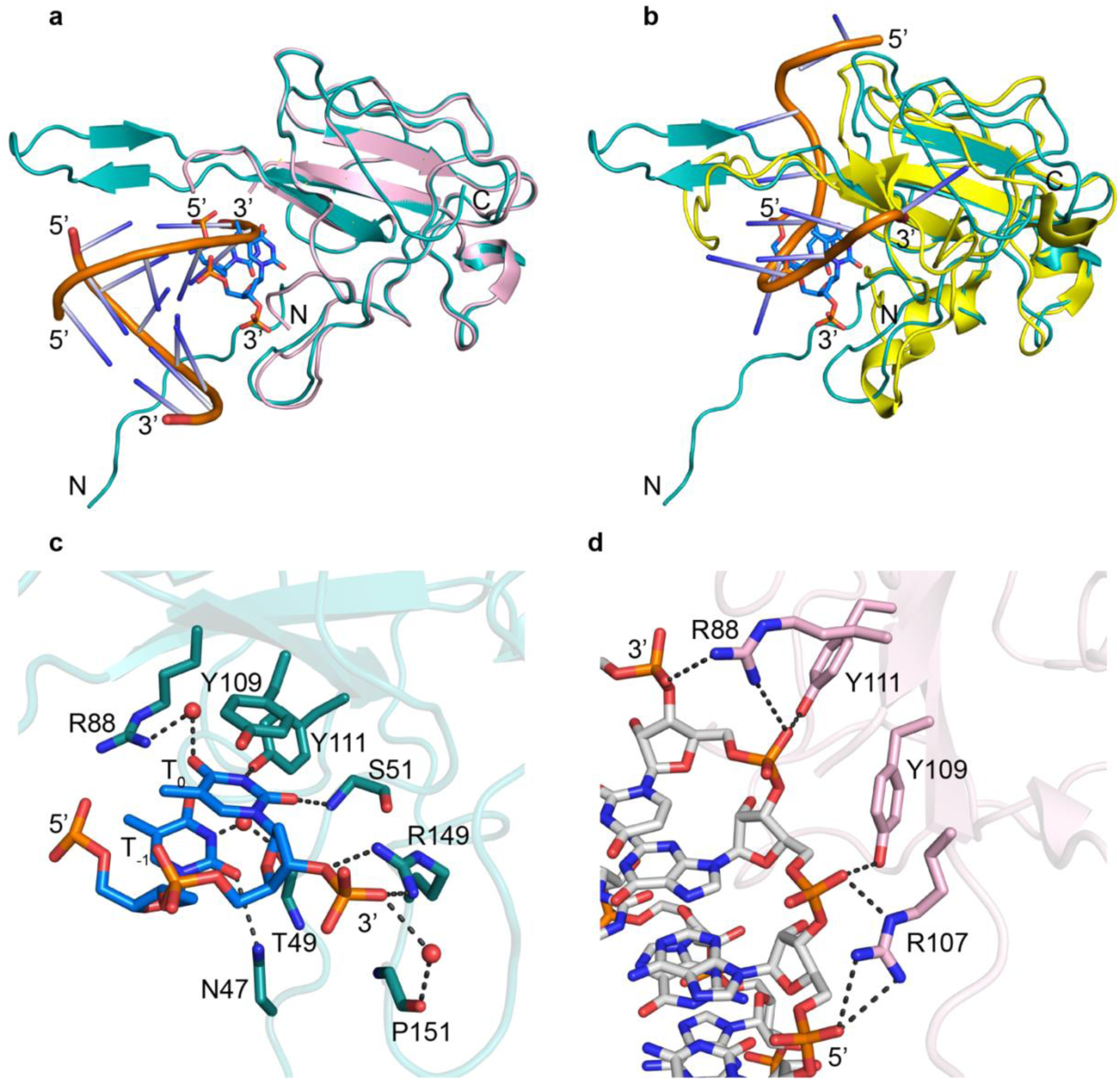
Interactions of N protein with nucleic acids. a) Superimposition of our structure (teal) with crystal structure of N-NTD complex with dsRNA (light pink, PDB ID: 7XWZ). b) Superimposition of our structure (teal) with NMR structure of N-NTD complex with ssRNA (yellow, PDB ID: 7ACT). RNA is shown as cartoon and ssDNA as stick. ssDNA binds to N-NTD at RNA binding pocket. c) Detail Interaction of ssDNA with N-NTD (teal, our structure). d) Interaction of dsRNA with N-NTD (light pink, PDB ID: 7XWZ). ssDNA and RNA are represented as sticks, water represented as red sphere. Gray dashed lines represent the hydrogen bonds. The bases of ssDNA interacts with protein through direct and water mediated hydrogen bonds. The RNA interacts with protein through hydrogen bonds in phosphate backbone.

## 3. Materials and Methods

### 3.1. Plasmid generation and protein purification

The protein expression vector was generated by cloning (cloning site: 5’ NdeI, 3’ BamHI) SARS-Cov-2 N gene (Gene ID: 43740575) in pET-11a vector without any affinity tag (GenScript). pET-11a expression vector carrying SARS-Cov-2 N gene was transformed in BL21 (DE3) cells (Invitrogen). Cells were grown in LB media at 37°C until reaching an optical density 0.5-0.6 at 600 nm. Then temperature was changed to 17°C and cells were induced by adding 0.2 mM IPTG at optical density 0.6-0.8. Cells were further grown overnight at 17°C. All the steps for protein purification were performed at 4°C unless specified.

E. coli cells (1 liter culture) were harvested by centrifugation and resuspended in 25 ml lysis buffer (20 mM Tris-Hcl, pH 7.5 + 200 mM Nacl + 1 mM DTT + 5% Glycerol) and protease inhibitor (Roche, Basel, Switzerland). The suspended cells were disrupted by sonication and then cell debris was separated by centrifugation at 12,000 rpm for 45 min.

The supernatant was treated with gradual addition of 5% polyethyleneimine to reach final polyethyleneimine concentration of 0.5% and stirred for 1 h to allow maximum precipitation. The precipitate was separated by centrifugation at 10 000 r.p.m. for 10 min. The supernatant containing the N-protein was saved.

The supernatant was added to SP Sepharose bead (6 ml, preequilibrated with lysis buffer) and agitated for overnight. N protein bound to SP Sepharose bead was passed through the column under gravity. After washing the bead by lysis buffer (40 ml), protein was eluted with 10 ml of elution buffer E1-E6 [lysis buffer having different concentration of salt (E1=0.3M, E2=0.4M, E3=0.5M, E4=0.6M, E5=0.8M and E6=1M NaCl)] gradually. The N protein was eluted at fraction E2 (0.4M NaCl) and E3 (0.5M NaCl) completely as confirmed by SDS-PAGE electrophoresis (Supplementary Fig. S2a). Eluted protein was further purified by Superdex-75 size exclusion column (GE Healthcare LifeScience) in FPLC buffer (20 mM Tris-Hcl, pH 7.5 + 100 mM Nacl + 1 mM DTT + 1% Glycerol) using an AKTA FPLC system (Supplementary Fig. S2b). Purity and concentration of the proteins were measured by gel electrophoresis and UV spectroscopy.

### 3.2. Mass Photometry

The mass photometry experiments were performed on mass photometer TwoMP (Refeyn Ltd., Oxford, UK). High precision cover glasses (coverslips) of size 24x50 mm of thickness No. 1.5H (Paul Marienfeld GmbH and Co KG, Lauda-Konigshofen, Germany) and six-well pre-cuts CultureWell gaskets (Grace Bio-Labs, Bend, Oregon, USA) were used. Coverslips were rinsed by deionized water and Isopropyl alcohol (IPA) in a sequence water – IPA – water – IPA – water and dried in a stream of ultrapure nitrogen. A drop of immersion oil Immersol 518F (Carl Zeiss Jena, Oberkochen, Germany) was placed on the microscope lens before placing coverslips on the mass photometer stage. Protein sample was prepared at 50nM and 100nM concentrations in sample buffer (20 mM Tris-Hcl, pH 7.5 + 100 mM Nacl + 1 mM DTT). The instrument was calibrated with a mix of beta-amylase from sweet potato (Sigma-Aldrich) and thyroglobulin from bovine thyroid (Sigma-Aldrich) dissolved in sample buffer. The autofocusing was performed with buffer free mode, then 15 μL of protein solution was placed on the coverslip well. The movies were recorded for 60 sec with regular view setting using AcquireMP (version 2023 R1.1) software. The experiments were triplicated with preparation of new probes. The data analysis was performed in histogram mode with manual selection of observed peaks and application of Gaussian fit using DiscoverMP (version 2023 R1.2) software.

### 3.3. Crystal Growth and Data Collection

We aimed to crystallize full length SARS-Cov-2 N protein in complex with ssDNA. Towards our aim, two different ssDNA (T-10=TTTTTTTTTT and T-20=TTTTTTTTTTTTTTTTTTTT) at 50% molar excess was mixed with purified SARS-Cov-2 N protein separately. Then the mixtures were concentrated to about 400 μM of protein concentration using Amicon Ultra-4 (Merck Millipore). Crystallization screening was performed using different commercially available crystallization screen by the sitting drop vapor-diffusion method at 4°C and 20°C. Crystal drops were set up by mixing 0.3 μl sample and 0.3 μl reservoir solution in a sitting drop 2-well crystallization plate (Molecular Dimension) using a robot, Mosquito Crystal (ttp Labtech). Crystals appeared after an extended period in a condition having 0.2M HEPES (pH 7.4) and 20% w/v PEG 4000 (PEG/pH, F-4 screen from Hampton Research). Crystals were cryoprotected using reservoir solution containing 20% v/v glycerol and flash frozen in liquid nitrogen. X-ray diffraction data were collected at the Southeast Regional Collaborative Access Team (SER-CAT) 22-ID beam line at the Advanced Photon Source, Argon National Laboratory.

### 3.4. Structure Determination and analysis

The collected diffraction data were indexed, integrated, and scaled using the HKL2000 program [75]. The crystals belong to the space-group I 41 with unit cell parameter (a=90.891, b=90.891, c=36.341, α=90, β=90, γ=90) (Table-1). Cell content analysis using Mathew’s coefficient [76] in CCP4 suggested that the unit cell of the crystal was not sufficient for full length SARS-Cov-2 N protein but able to accommodate half of the full-length protein. We determined the phase by molecular replacement using the program Phaser [77]. Previous structures of N-NTD (PDB ID:7N0R) and N-CTD (PDB ID:6ZCO) together or N-CTD (PDB ID:6ZCO) alone used as search model did not give any solution. However, structures of N-NTD (PDB ID:7N0R) used as search model gave a solution successfully. The structure was solved at 1.55Å resolution by molecular replacement, using the program Phaser [77] and N-NTD (PDB ID:7N0R) structure as search model. Model building of the protein and bound DNA were manually performed using the program Coot [78]. The structure model was refined by Phenix refinement [79,80] and final model was validated with PDB validation tool and Molprobity [81]. Structural figures were created using the software PyMOL. Data collection statistics and refinement parameters of the crystal structure is given in table-1.

## 4. Conclusion

In this study, we successfully purified the SARS-CoV-2 N protein using an optimized protocol that yielded highly soluble, homogeneous, nucleic acid-free protein suitable for structural studies. Our purification method included polyethyleneimine precipitation, SP Sepharose chromatography, and size exclusion chromatography, resulting in the isolation of N protein with significant purity and confirmation of its dimeric form via mass photometry. These findings aligned with previous studies that demonstrated the N protein’s tendency to dimerize under similar conditions. Our purification method provides a reliable source of N protein for further exploration of its biochemical and biophysical properties, specifically regarding its interactions with nucleic acids.

Despite our efforts to crystallize the full-length N protein in complex with ssDNA, we encountered the inherent challenges posed by its flexibility and protease sensitivity, leading to the crystallization of only the N-terminal domain (NTD). Our structural analysis revealed that the NTD, along with a part of the flexible N-arm, formed a specific arrangement, potentially contributing to the assembly of the viral ribonucleoprotein complex (RNP). Notably, the crystallized NTD displayed a monomeric form in complex with a trinucleotide, suggesting a distinct mode of interaction with ssDNA compared to RNA. These structural insights, combined with previous findings, expand our understanding of the N protein’s role in viral assembly and its potential as a target for nucleic acid-based therapeutics.

Finally, our results demonstrate the importance of the N-arm in the N protein’s interaction with nucleic acids, potentially playing a role in RNP formation. The unique binding mode of ssDNA observed in our structure highlights key residues involved in this interaction, providing a foundation for future therapeutic development. Given the challenges of crystallizing the full-length N protein, further experiments are required to fully elucidate its role in RNP assembly and its interactions with different nucleic acids. These findings open new avenues for designing inhibitors targeting the N protein, with the potential to disrupt SARS-CoV-2 replication and pathogenesis.

## Author contributions

A.M. designed and performed all the experiments, structural analysis and interpretation; A.M. prepared all the figures; A.M. and H.M. wrote the paper; A.M. conceived the project and H.M. secured funding.

## Data Availability Statement

Atomic coordinates and structure factors have been deposited in the Protein Data Bank with accession codes 9CJ6 (https://www.rcsb.org/structure/9CJ6).

Data are available from A.M. and H.M upon request

## Acknowledgements

The authors would like to acknowledge Dr. Sergey G. Tarasov and Ms. Marzena A. Dyba at the Biophysics Resources of CCR/NIH for providing instruments and technical support.

## Supplementary Materials

**Supplementary Figure S1:**
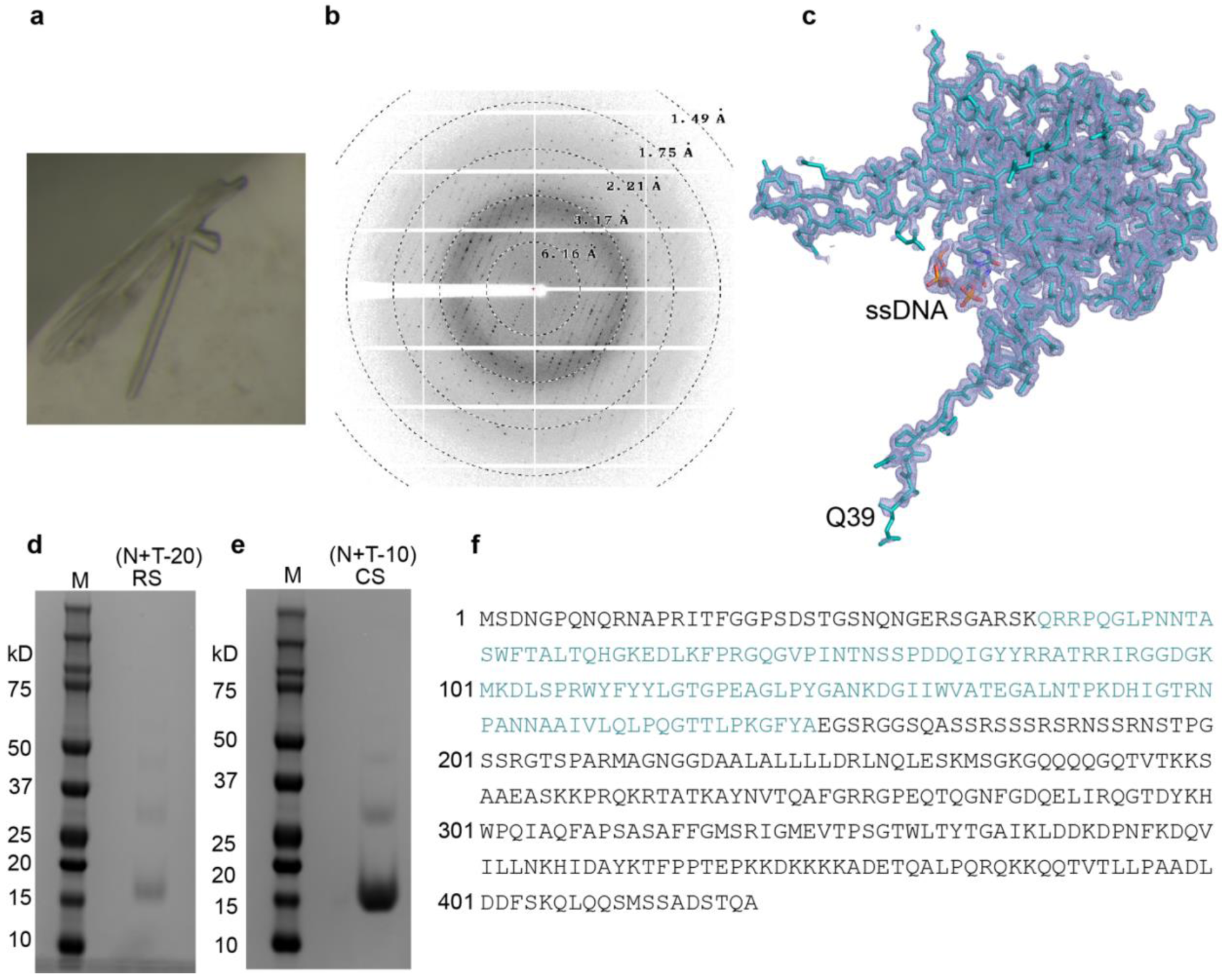
Crystallization of SARS-CoV-2 N protein with Poly-T (T-20) ssDNA. a) Crystal appeared in a condition after an extended period. b) Cryoprotected crystal diffracted at high resolution. c) The crystal appeared in the I 41 space group with unit cell parameter (a=90.891, b=90.891, c=36.341, α=90, β=90, γ=90). The cell content analysis suggested that the unit cell volume can accommodate half of the full-length N protein. Molecular replacement with only N-NTD (PDB ID:7N0R) successfully provided a solution, resulting a single protein molecule per asymmetric unit. d & e) The SDS-PAGE analysis of residual samples (RS; N+T-20)) and (CS; N+T-10) from crystallization wells confirmed that the full-length protein was digested under the crystallization condition. c & f) We modeled residues Q39 to A173 (blue sequence) of full-length N (black and blue sequence) protein in complex with ssDNA.

**Supplementary Figure S2:**
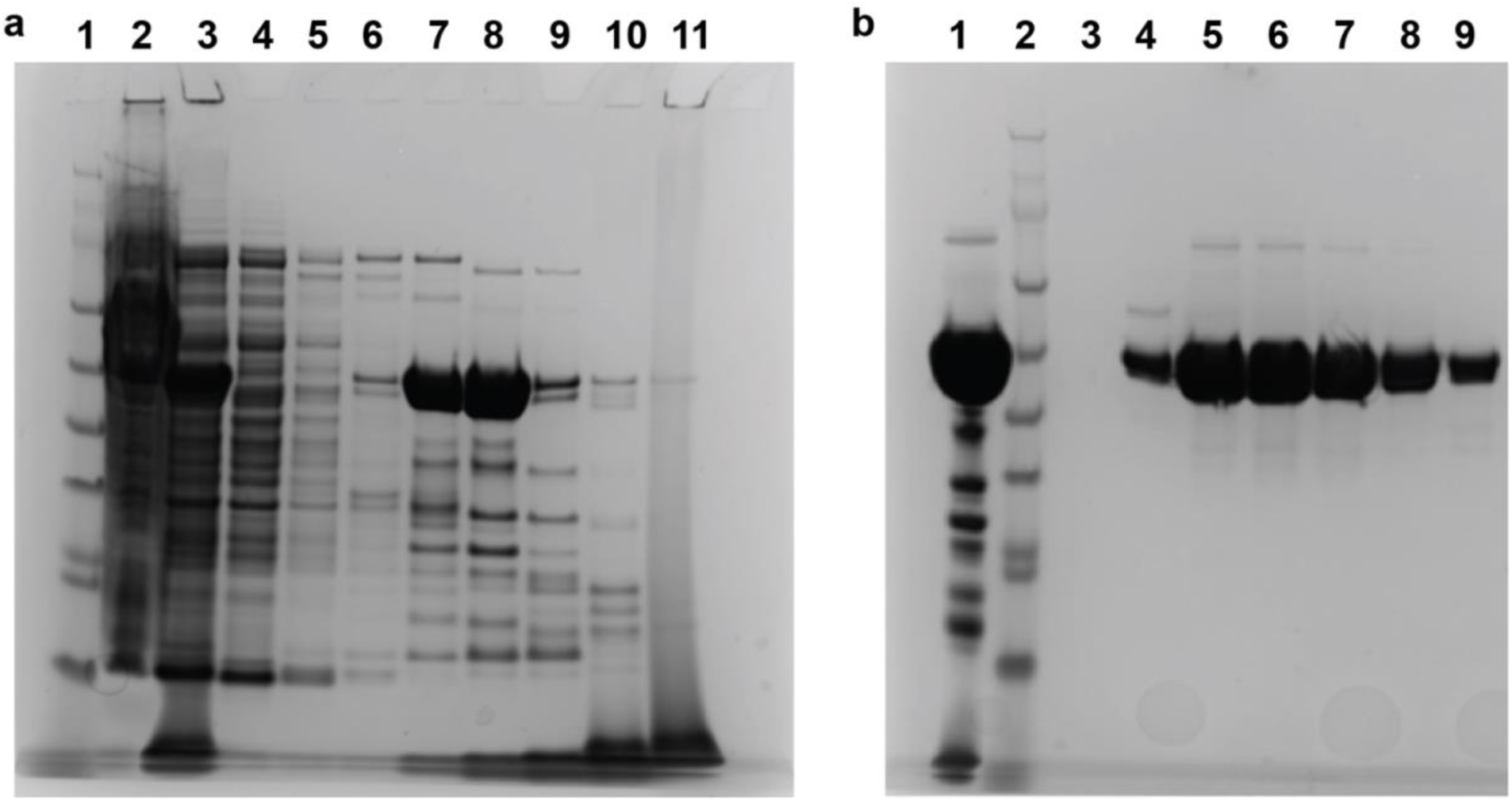
SDS-PAGE analysis of SARS-CoV-2 N protein expression and purification. a) Lane-1: NuPAGE protein standard marker; lane-2: Supernatant after sonication; lane-3: Supernatant after polyethyleneimine precipitation; lane-4: SP-column flow through; lane-5: SP-column wash; lane-6: Elution 1 (E1=0.3M NaCl); lane-7: Elution 2 (E2=0.4m NaCl); lane-8: Elution 3 (E3=0.5M NaCl); lane-9: Elution 4 (E4=0.6M NaCl); lane-10: Elution 5 (E5=0.8M NaCl); lane-11: Elution 6 (E6=1M NaCl). b) SDS-PAGE of Superdex-75 size exclusion chromatography. Lane-1: injected sample, lane-2: NuPAGE protein standard marker, Lane-3-9: FPLC fractions 14-20.

